# Phylogram instead of chronogram when assessing the neutral evolution of a trait

**DOI:** 10.1101/2025.06.16.659089

**Authors:** T. Latrille, T. Gaboriau, N. Salamin

## Abstract

Comparative analyses of trait evolution aim to uncover the different evolutionary forces shaping phenotypic diversity among species. This is typically done by fitting various evolutionary models to observed trait changes along a species tree. For instance, under neutral evolution, trait values are modelled as changing randomly along the branches of the tree. In contrast, for a trait under selection, species are typically assumed to track an optimal trait value, which itself may shift along the tree. Here, rather than relying on alternative models to discriminate among evolutionary scenarios, we focus on the underlying species tree, and specifically the units in which its branch lengths are measured. Species trees are usually represented as chronograms, with branch lengths proportional to time. The rationale is that time correlates with trait changes through its connection to the number of generations. However, since the generation time of species can also vary along the phylogenetic tree, we argue that chronograms introduces biases. In contrast, phylograms with branch lengths in units of sequence divergence will account for the effect of changing generation time. In this study, we develop a method to test whether a phylogram provides a better fit than a chronogram for modelling trait evolution. We show using simulations that, for a trait evolving neutrally, the fit of a random evolution model has more support on a phylogram than on a chronogram. However, comparing models and testing different scenarios of selection using a phylogram leads to incorrect predictions. Given these results, we argue that we should use phylograms instead of chronograms when assessing the neutral evolution of a trait. Nevertheless, we support the fact that we should generally continue to use chronograms to model selection acting on a quantitative trait.

**Lay summary:** When we look at different species, we can see that some traits are different between them. For example, the brain size of different primate and human lineages varies. The question we ask is: does the time that passed explain the differences in their traits? Or are genetic differences between species better at explaining the differences? The standard in evolutionary biology is to use time to explain differences in traits. However, if the trait is not under natural selection and instead is changing due to accumulating mutations, the genetic differences between species instead of time should best explain the differences in traits. As a result, we argue that using both genetic differences and species divergence time simultaneously provides a better understanding of how traits evolve across species.

## Introduction

By measuring changes in a phenotypic trait along a lineage, it is possible to infer the type of selection acting on it or whether this change is only due to random genetic drift. At a larger scale, from the observed variations of traits across species, regimes of evolution are typically assessed using phylogenetic comparative methods, where traits are modelled as evolving along the branches of the species tree (Felsenstein, 1985; Felsenstein, 1988; Harmon, 2018). For example, to model neutral evolution, the mean trait value is said to follow a Brownian motion (BM), branching and evolving independently after each speciation event (Felsenstein, 1985; Lynch and Hill, 1986; Hansen and Martins, 1996). In other words, for each branch of the tree, the value at the descendant node is normally distributed around the ancestral value, with a variance proportional to the branch length. In this framework, reconstructing trait variation along the whole phylogeny as a BM can thus constitute a null model of neutral trait evolution.

Alternatively to a simple BM, adding a trend parameter in the BM is interpreted as a signature of directional selection at the phylogenetic scale (Silvestro et al., 2019). More complex models to detect selection have been proposed, notably the Ornstein-Uhlenbeck (OU) processes, where trait variation is constrained around an optimum value, which is often interpreted as a signature of stabilizing selection (Hansen, 1997; Butler and King, 2004; Beaulieu et al., 2012). Methodologically, this alternative model of evolution raises issues since an OU process might be statistically preferred over a BM due to sampling artifacts (Silvestro et al., 2015; Cooper et al., 2016; Price et al., 2022). Adding biological realism, the OU process can be relaxed to allow for multiple optima along the phylogenetic tree, which is interpreted as a few abrupt changes in the environment along the phylogeny (Ingram and Mahler, 2013; Uyeda and Harmon, 2014; Khabbazian et al., 2016; Mitov et al., 2020; Grabowski et al., 2023). Finally, the optimum can also be allowed to change continuously along the tree, which results in the trait itself reflecting the movement of the optimum along the lineages due to the constant adaptive evolution of the trait toward the optimum (Hansen and Martins, 1996; Hansen, 2024). One special case of a continuously changing optimum is again the BM, where changes in the optimum are a consequence of the randomly changing environment (Hansen et al., 2008), named fluctuating selection (Holstad et al., 2024) or also diversifying selection due to the diversity of generated phenotypes at the clade level (Latrille et al., 2024). This modelling raises an issue in disentangling selection from neutral evolution: the best fit of BM can not be interpreted as the trait evolving neutrally, it can also be interpreted as a continuously moving optimum.

In order to disentangle neutral evolution from selection, another approach is to contrast the observed rate of evolution to the neutral expectation (Lande, 1980). The neutral expectation can be obtained if the underlying genetic architecture of the trait is known and the trait encoded by many loci of additive effects (Barton et al., 2017; Sella and Barton, 2019). Additionally, we can also derive the expected changes in mean trait value along a lineage given the trait variance at the population scale (Turelli, 1984; Felsenstein, 1988). When generalized to many species at the phylogenetic scale, contrasting between and within-species variations allow us to disentangle neutral evolution from selection (Latrille et al., 2024). More generally, comparing the rate of evolution at different timescales allows inferring the regime of evolution (Hansen, 2024; Holstad et al., 2024). However, trait variation at different timescales is not always available, and the genetic architecture of the trait is often unknown. Instead, to disentangle neutral evolution from selection in the context of phylogenetic comparative methods, we hereby focus on the underlying tree and the unit of its branch lengths (Fig. 1A).

**Figure 1:**
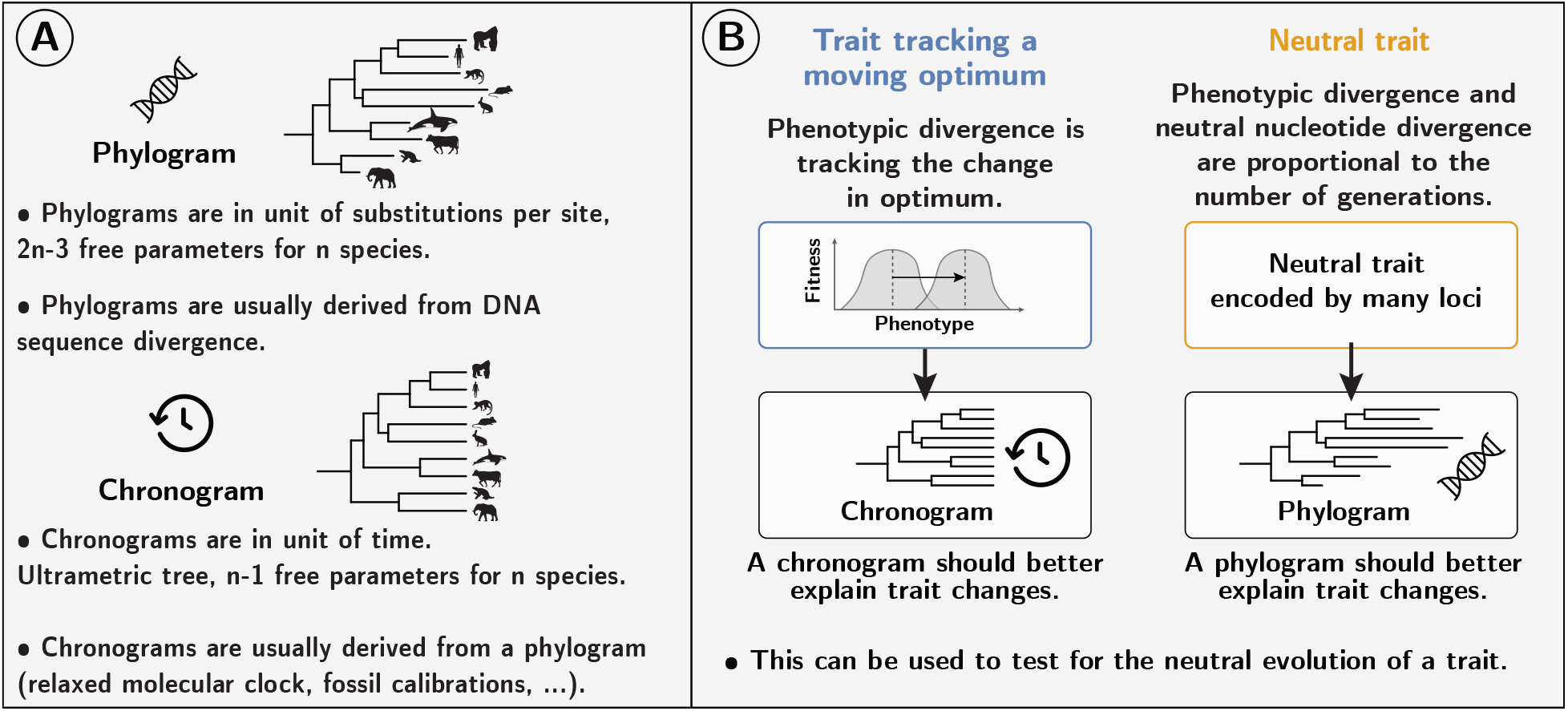
Panel A: Across many species, the evolution of continuous traits can be modelled as a stochastic process evolving along the branches of a phylogenetic tree. The branches of such phylogenetic tree can be measured in either time (chronogram) or number of substitutions (phylogram). Panel B: Theoretically, changes of a trait under a moving optimum should be better predicted by a chronogram. Instead, changes of a neutral trait should be better predicted by a phylogram.

Typically, the tree is assumed to be known and obtained independently, and the *de facto* standard in phylogenetic comparative methods is to use a chronogram, where the branch lengths are proportional to time (Felsenstein, 1985; Harmon, 2018). Theoretically, for a neutrally evolving trait, trait changes depend directly on the number of generations (Hansen and Martins, 1996), which is proportional to time only if the average time between two consecutive generations, called generation time, is constant and equal for all species. Since generation time depends on the time for an individual to reach sexual maturity, it can vary considerably between species, from a few months for coral reef pygmy gobies to 150 years for the Greenland shark (De Magalhães and Costa, 2009; Nielsen et al., 2016). As a result, variations of generation time between species means that modelling neutral evolution as a BM on a chronogram might induce biases (Litsios and Salamin, 2012).

Alternatively, on a phylogram, the branch lengths represent another quantity than time, which can be used as the backbone to model trait evolution and can potentially absorb changes in generation time. For example, it is known that using phylograms in units of morphological distance can be used to improve ancestral trait reconstruction for a discrete character (Wilson et al., 2022). From a genomic perspective, phylograms can also be in units of nucleotide divergence, that is, depicting the number of nucleotide substitutions occurring along a branch. Because the number of neutral nucleotide substitutions accumulating along a branch is proportional to the number of generations (Kimura, 1968; Kimura, 1983), nucleotide divergence would in this case absorb the effect of changing generation time (i.e. longer branches for lineages with shorter generation time). Such a phylogram, if obtained from neutrally evolving genomic loci would be more appropriate to model a trait evolving neutrally (Latrille et al., 2024). Additionally, the rational to use nucleotide divergence to absorb changes in generation time is also valid for changes in mutation rate (per loci and per generation) under the assumption of a constant genetic architecture of the trait and that changes in mutation rate impact the whole genome (Latrille et al., 2024). Empirically, such changes in mutation rate along a phylogenetic tree are also observed, although to a lesser extent than changes in generation time (Bergeron et al., 2023).

Under the alternative scenario involving selection, shorter generation time and higher mutation rate can also allow for the species to track the changing optimum faster and not lag behind (Lande, 1976; Hansen et al., 2008). However, the timescale for this lag (hundreds or thousands of years) is negligible compared to the timescale on which the optimum changes at the level of clades (millions of years) in a first approximation (Hansen, 2024). Thus, at the phylogenetic scale, since shifts in the optimum fitness peak is extrinsic to the species and is dependent on the environment which varies with time, using a chronogram would be more accurate to model mean trait changes. As a result, for a trait under selection, mean trait changes should be better explained by the time rather than the number of substitutions that occurred along a branch (Fig. 1B).

In this study, we test this hypothesis, namely if using a phylogram (in units of nucleotide divergence for neutral loci) instead of a chronogram (in units proportional to time) can allow to assess the regime of evolution of a trait, by evaluating the fit of a BM on both trees. More precisely, we seek to test if the changes of a neutral trait are better predicted by a phylogram. Conversely, we also seek to test if changes of a trait under a moving optimum are better predicted by a chronogram. Using genomic information we seek to provide additional approaches to model trait evolution, particularly to disentangle selection and neutral evolution.

## Methods

To assess whether a phylogram is better suited to model neutral evolution we first performed simulations of a trait evolving along a phylogenetic tree under different regimes of selection (neutral, moving optimum, multiple optima). Second, we fitted a single rate Brownian motion (BM) to the simulated data, and we compared the fit of the BM on a phylogram versus a chronogram. Third, based on the single rate BM, we derived a model to estimate the support of a phylogram over a chronogram that we apply to the simulated dataset. Fourth, we fitted several models of trait evolution to the simulated datasets: a multi-rate BM, an Ornstein-Uhlenbeck (OU) process, and a relaxed OU with multiple optima, tested on a phylogram versus a chronogram. Finally, we gathered an empirical dataset of body and brain masses from mammals, including both a chronogram and a phylogram (in units of nucleotide divergence from neutral loci), on which we tested the support of a phylogram for the single rate BM.

### Simulations along a phylogenetic tree

We performed simulations under different selective regimes (neutral, moving optimum, multiple optima). Simulations were individual-based and followed a Wright-Fisher model with mutation, selection and drift for a diploid population including speciation along a predefined ultrametric phylogenetic tree. We used the same simulation framework as in Latrille et al. (2024), with parameters detailed in the supplementary material (section 1). The parameters of simulations were chosen to mimic an empirical dataset of mammals. In summary, the trait was encoded by *L* independent loci, with each locus contributing additively, and mutations were drawn from a Poisson distribution at each generation. Parents were selected for reproduction according to their phenotypic value, with a probability proportional to their fitness. Flattening the fitness landscape resulted in neutral evolution, which meant that each individual had the same probability of being sampled at each generation regardless of its trait value. Alternatively, for a trait under selection around a moving optimum, we modelled stabilizing selection acting on the trait, with the optimum value changing as a geometric Brownian motion (BM) along the phylogenetic tree (Hansen, 1997; Hansen and Martins, 1996). Finally, for a trait under selection around multiple optima (Uyeda and Harmon, 2014), we draw a Bernoulli variable to test whether there is a switch of optimum along a branch; if a switch occurs, the sliding of the optimum is drawn from a reflected exponential distribution (symmetric positive and negative values).

At each internal node of the tree (i.e. speciation event), the population is split into two daughter populations running independently on each of the two branches, and the process is repeated until the tips of the tree are reached. As a control, we performed simulations with constant mutation rates, generation times and effective population sizes (*N*_e_). Alternatively, we performed simulations with fluctuating mutation rates, generation times and *N*_e_, where we used a BM to model the long-term changes along the phylogenetic tree, and we overlaid short-term changes in *N*_e_ (see supplementary material section 1).

The phylogram is obtained from the same simulation setting on a set of 30,000 independent neutral loci, and the nucleotide divergence is computed as the number of substitutions along each branch. The chronogram is derived from the phylogram by fitting a relaxed molecular clock model (correlated rate model with penalized likelihood, default value) to the phylogram with the *R* package *ape* (Paradis et al., 2004).

### Brownian motion (BM)

On the simulated dataset, we fitted a single rate BM using either a phylogram or a chronogram as the underlying tree. We used *RevBayes* (Höhna et al., 2016) to fit a Brownian motion (BM) on continuous traits evolving along the branches of a phylogenetic tree (Felsenstein, 1985; Felsenstein, 1988). The data consist of the mean trait value for each extant species, and the tree topology is fixed. Along each branch, the value of the trait is drawn from a normal distribution with mean equal to the parent node value and variance equal to the branch length (Felsenstein, 1985). Formally, the BM process is described by the equation:

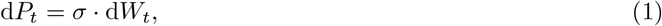

where *P*_*t*_ is the trait value at time *t, σ* is the rate of the BM, and d*W*_*t*_ is a Wiener process (i.e. a standard Brownian motion). We used a log-uniform prior for *σ*. Using Markov chain Monte Carlo (MCMC), we reconstructed the ancestral trait value at each node of the tree as the posterior mean estimate (burn-in of 1000 gen., running of 10,000 gen., 2 chains), and we compared the accuracy of the reconstruction using either a phylogram or a chronogram as the underlying tree. Both trees are scaled such that the sum of all the branch lengths is equal to one in each case. Importantly, the data and the tree topology are the same in both analyses; only the branch lengths are different. In this simulation setting, for traits simulated under neutral evolution, we expect more accurate trait reconstruction on a phylogram, and conversely, for traits under selection (moving optimum), more accurate trait reconstruction on a chronogram.

### BM with a switch

To test the support of a phylogram over a chronogram, we implemented a model based on a single rate BM. The model was implemented in *RevBayes* and contains a switch variable, denoted as *π* that allows the model to switch the branch lengths in units of time to units of substitutions (Fig. 2A). Formally, *π* ∈ 0, 1 is a Bernoulli random variable (*π* ∈ 0, 1) with a prior probability of 0.5. If *π* is 0, the branch lengths are those of the chronogram, and if *π* is 1, the branch lengths are those of the phylogram. The tree topology is fixed, meaning both trees have the same branching structure, but because branch lengths are different and not necessarily on the same scale, both trees are re-scaled by dividing branch lengths by the total tree length. As in the previous section, a BM is fitted to the data by modelling trait changes as normal distributions (*dnNormal* in *RevBayes*) running along the branches of the tree, with variance proportional to the branch length and the squared value of the Brownian rate parameter (*σ*, log-uniform prior). Alternatively for large trees or for faster computation, the likelihood can also be estimated by the REML method (*dnPhyloBrownianREML* in *RevBayes*). Altogether, the input data is the mean trait value for extant species and both the chronogram and phylogram with the same tree topology. The posterior mean of *π* (burn-in of 1,000 gen., running of 10,000 gen., 2 chains) is the probability that the phylogram is favoured over the chronogram. We expect *π* = 1 for a trait under neutral evolution, and *π* = 0 for a trait under selection (moving optimum).

**Figure 2:**
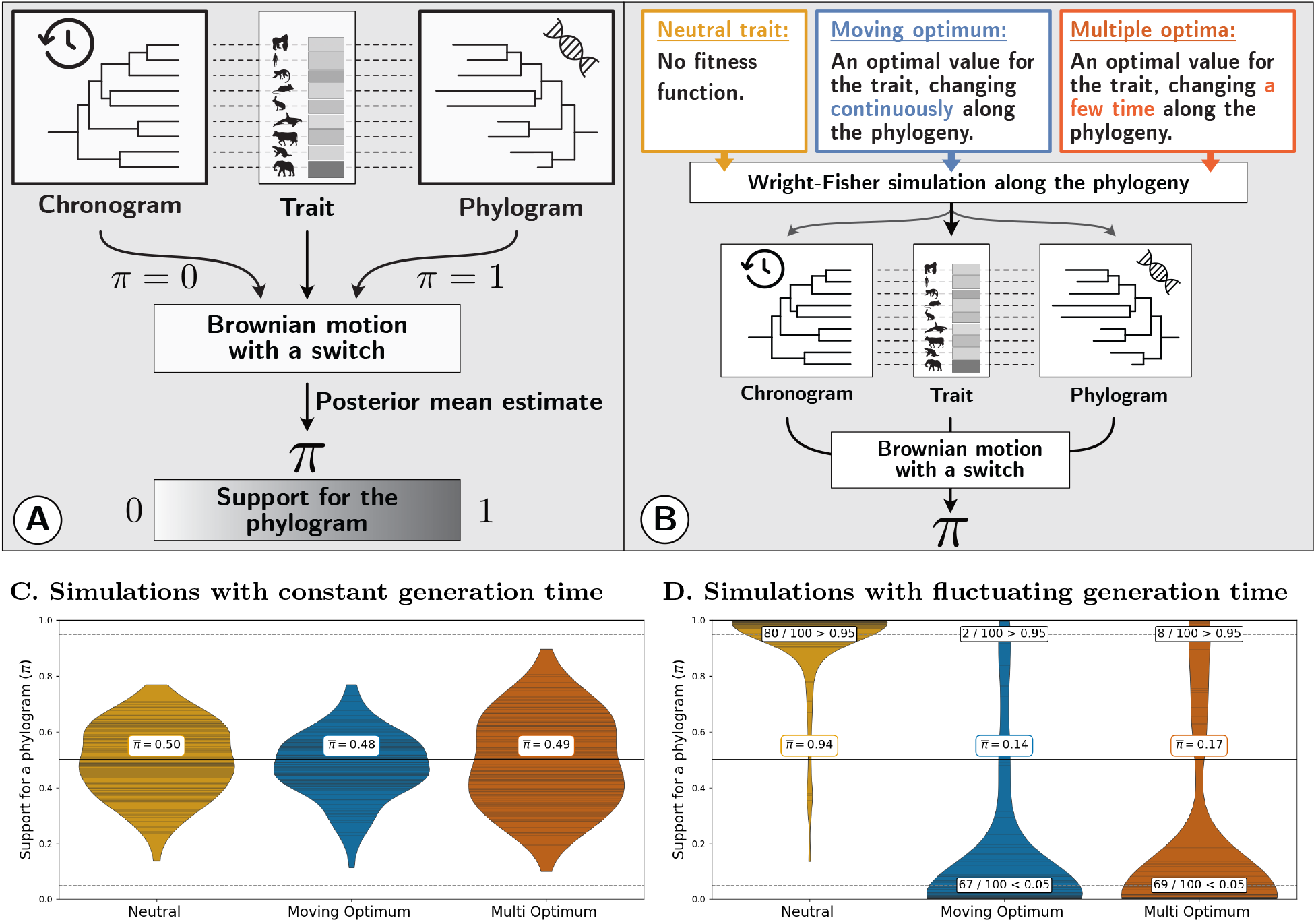
Testing the support for a phylogram over a chronogram. Panel A: Brownian motion (BM) is used to model trait changes along a phylogeny, given a mean trait value for extant species. The model contains a switch variable, denoted *π*, that switches the branch lengths in units of time (*π* = 0, chronogram) to units of nucleotide divergence for neutral sites (*π* = 1, phylogram). The posterior mean estimate of *π* indicates whether a BM supports better the phylogram over the chronogram (0 ≤ *π* ≤ 1). Panel B: Wright-Fisher simulations of trait evolution along a phylogeny. At each node of the tree a speciation event creates two descendant species evolving independently. Neutral evolution is modelled as a constant fitness regardless of the phenotype of individuals (yellow). Selection is modelled by stabilizing selection around a optimum value, which is changing along the phylogenetic tree: as a moving optimum (blue) or as multiple optima (red). The output of the simulation is: 1) the mean trait value for each extant species, 2) the phylogram as obtained from independent neutral markers and 3) the chronogram derived from the phylogram assuming a relaxed molecular clock. The posterior mean estimate of *π* is inferred from the output of the simulation. Panels C & D: Violin plot of the posterior mean of *π* across 100 replicates for the different simulated regimes of evolution with mean *π* values at the center 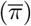. Horizontal lines inside the violins are the estimated *π* of each replicate simulations. Results of simulations with constant generation time (Panel C) and with fluctuating generation time, mutation rate and effective population size (Panel D).

### Alternative models of evolution

In addition to a single rate BM, we also fitted a BM with multiple rate parameters (Eastman et al., 2011), using either a phylogram or a chronogram as the underlying tree to the simulated dataset. The likelihood of the data is estimated by the REML method (*dnPhyloBrownianREML* in *RevBayes*), and the number of rate shifts is estimated by reversible-jump MCMC (*dnReversibleJumpMixture* in *RevBayes*). For each branch, we draw a rate-multiplier either equal to 1 (no rate shift), or drawn from a log-normal distribution with a median of 1, and a standard deviation such that rate shifts range over about one order of magnitude. We set our prior to the expected number of rate shifts of 1 across the whole tree. The number of rate shifts as well as the variance of rate parameters are estimated as the posterior mean (burn-in of 1,000 gen., running of 50,000 gen., 2 chains).

Also, we compared the fit of an Ornstein-Uhlenbeck (OU) process (eq. 2) to a BM (Hansen, 1997; Butler and King, 2004), using either a phylogram or a chronogram as the underlying tree. The OU process is described by the equation:

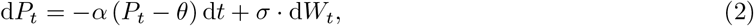

where the parameter *α* in eq. 2 is the strength of the pull towards the optimum, *θ* is the optimum value and the other parameters are defined as in eq 1. The likelihood of the data is estimated by the REML method (*dnPhyloOrnsteinUhlenbeckREML* in *RevBayes*), and because the OU process has more parameters than the BM, we used a reversible-jump MCMC switch between the two models (0 for BM and 1 for OU, *dnReversibleJumpMixture* in *RevBayes*). The prior for *θ* is uniform between the minimum and maximum trait value for extant species. The prior for *α* is a reversible-jump mixture distribution: a value of 0 or drawn from an exponential distribution with mean equal to half the root age divided by ln(2), meaning that we expect a phylogenetic half life of half the tree age. Thus, when *α* = 0 the OU process (eq. 2) is equivalent to a BM (eq. 1). The support for the OU model is estimated as the posterior mean of the switch variable (*p*_OU_), or equivalently that of *α*≠ 0 (burn-in of 1,000 gen., running of 10,000 gen., 2 chains).

Finally, we fitted a relaxed OU with multiple optima (Uyeda and Harmon, 2014), using either a phylogram or a chronogram as the underlying tree. The likelihood of the data given the tree is estimated by the REML method (*dnPhyloOrnsteinUhlenbeckREML* in *RevBayes*), and the number of optimums is also estimated by the reversible-jump MCMC method (*dnReversibleJumpMixture* in *RevBayes*) as the posterior mean estimate (burn-in of 1,000 gen., running of 5,000 gen., 2 chains). For each branch, we draw a shift in optimum either equal to 0 (no shift), or drawn from a uniform distribution centred on 0 with breadth spanning the whole range of observed trait value. We set our prior to the expected number of optimum shifts of 1 across the whole tree.

### Empirical dataset

We analysed a dataset of body and brain masses from mammals. The log-transformed values of body and brain masses were taken from Tsuboi et al. (2018). We removed individuals not marked as adults and split the data into males and females due to sexual dimorphism in body and brain masses. We also extracted brain and body masses from the *COMBINE* dataset (Soria et al., 2021). The mammalian genomic data are gathered from the Zoonomia project (Genereux et al., 2020). More specifically, the phylogram in units of nucleotide divergence is estimated on a set of neutral markers in Foley et al. (2023). The chronogram is derived from the phylogram by fitting a relaxed molecular clock model (correlated rate model with penalized likelihood, default value) to the phylogram with the *R* package *ape* (Paradis et al., 2004).

## Results

### Support for a phylogram over a chronogram

We tested the hypothesis that a phylogram is better suited to model neutral evolution than a chronogram. We simulated traits evolving on a phylogenetic tree under different regimes of selection: 1) neutrally evolving, 2) under a moving optimum, and 3) under multiple optima (see Methods). We first assessed the accuracy of ancestral trait reconstruction on a phylogram versus a chronogram, under the assumption that changes follow a BM. For a neutral trait, ancestral trait reconstruction was more accurate on a phylogram (Fig. S2A, *r*^2^ = 0.95) than on a chronogram (Fig. S2B, *r*^2^ = 0.85). In contrast, for a trait evolving under a moving optimum, ancestral trait reconstruction was less accurate on a phylogram (Fig. S2C, *r*^2^ = 0.79) than on a chronogram (Fig. S2D, *r*^2^ = 0.96).

We then tested for the BM support of phylogram over a chronogram, and we developed a statistic denoted as *π* (Fig. 2A), which ranges from 0 (full support for the chronogram) to 1 (full support for the phylogram). For simulations with a constant generation time, the support is similar for both a phylogram and a chronogram regardless of the regime of evolution (Fig. 2B-C), which is expected since the branch lengths are equivalent in both trees. Next, we simulated changing generation time, mutation rate (per generation) and effective population size (*N*_e_) along the phylogenetic tree, mimicking a mammalian range of changes. In this case, for a simulated neutral trait, the BM was better fitted on a phylogram than on a chronogram, with an average support for phylogram of 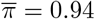 across 100 replicate simulations, and 80 of the 100 replicates above the 0.95 threshold (Fig. 2D, yellow). Conversely, for a simulated trait under selection, the fit of a BM on a chronogram was better than on a phylogram with an average support for phylogram of 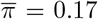 for multiple optima (69 out of 100 replicates below the 0.05 threshold, Fig. 2D, blue) and 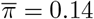 for a moving optimum (67 of the 100 below the 0.05 threshold, Fig. 2D, red).

On the empirical mammalian dataset from Tsuboi et al. (2018), for body mass the support for the phylogram is *π* = 0.0, when sex is not taken into account. When splitting the dataset into males and females, the support for the phylogram is *π*_♀_ = 0.85 and *π*_♂_ = 0.70. For brain mass, the support for the phylogram is *π* = 0.0035 on the mixed dataset, with *π*_♀_ = 0.56 and *π*_♂_ = 0.51. On the *COMBINE* dataset, which does not distinguish between males and females, the support for the phylogram is *π* = 0.0 for body mass and *π* = 0.0015 for brain mass (Table S1). We also computed *π* using an approximate likelihood computation (REML), a method that allow for faster computation and scale better for larger trees, and found similar results (Table S1).

### Fitting alternative models of evolution

Besides testing the fit of a BM with a single rate, we also fitted a multi-rate BM (see Methods). When fitting a multi-rate BM, the estimated variance of rate parameters (*v*) was lower on a phylogram than on a chronogram for a trait evolving neutrally (Fig. 3A, yellow violins, Wilcoxon paired test with *p*_value_ = 8.8 *×* 10^−14^), and the number of rate shifts (*n*) was reflecting the prior on the phylogram while not on the chronogram, showing a bias (Fig. 3B, yellow violins). Conversely, for a trait under a moving optimum, *v* was higher on a phylogram than on a chronogram (Fig. 3A, blue violins, Wilcoxon paired test with *p*_value_ = 8.9 *×* 10^−15^), and *n* was reflecting the prior on the chronogram while not on the phylogram, showing a bias (Fig. 3B, blue violins). Altogether, fitting a BM on a phylogram is more accurate for a trait evolving neutrally, but results in less accurate estimates for a trait under selection.

**Figure 3:**
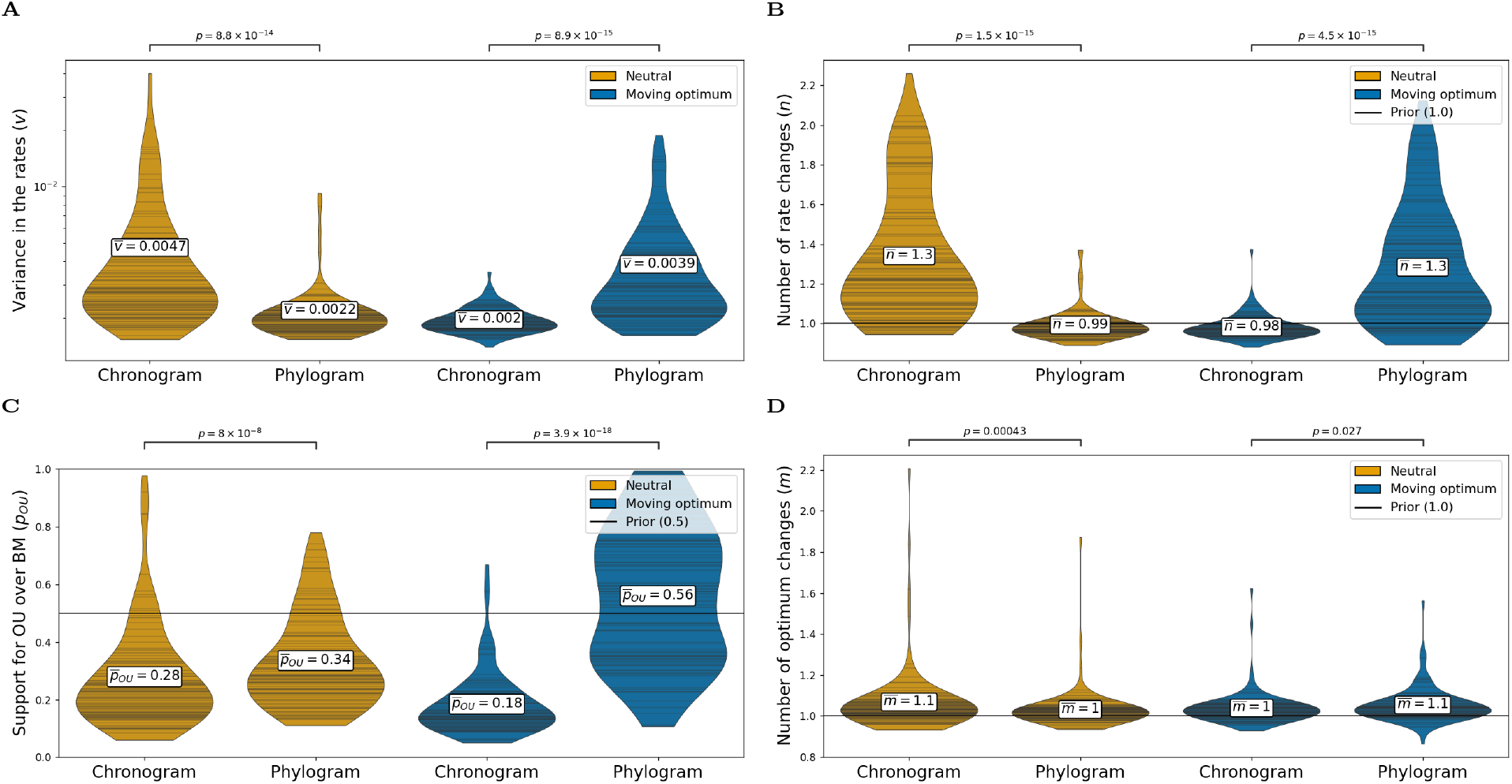
Mis-classification of trait evolution on a phylogram. Violin plot of posterior parameter estimates for different models of trait evolution and different simulated regimes of selection. Horizontal lines inside the violin show the results of each replicate simulation. Simulations of 100 replicates per regime: trait evolving under a neutral regime (yellow) and under a moving optimum (blue). Wilcoxon rank-test are performed between the paired estimates on the phylogram and the chronogram. Panel A & B: Fit of a relaxed Brownian motion (BM) with multiple rate parameters, using either a phylogram or a chronogram. Posterior estimate for the variance in rate parameters (Panel A) and number of rate changes (Panel B). Panel C: Relative fit of an Ornstein-Uhlenbeck (OU) process compared to BM, using either a phylogram or a chronogram. Posterior estimates for the support of the OU process over the BM (reversible-jump). Panel D: Fit of a relaxed OU with multiple optima, using either a phylogram or a chronogram. Posterior estimate for the number of optimum changes.

Moreover, we tested alternatives to the BM with OU models: with a single optimum and with multiple optima (see Methods). First, for a trait under a moving optimum, the estimated support for the OU process (*p*_OU_) was higher on a phylogram than on a chronogram (Fig. 3C, blue violins, Wilcoxon paired test with *p*_value_ = 3.9 *×* 10^−18^), an expected result since the BM with a switch supported the chronogram. For a trait evolving neutrally, *p*_OU_ was still higher on a phylogram than on a chronogram (Fig. 3C, yellow violins, Wilcoxon paired test with *p*_value_ = 8.0 *×* 10^−8^), an unexpected result since the BM with a switch supported the phylogram, thus showing a high rate of mis-classification on a phylogram. When fitting a multi-OU process on a trait evolving neutrally, the number of optimum shifts (*m*) was reflecting the prior on the phylogram, but not on the chronogram (Fig. 3B, yellow violins). Conversely, for a trait under a moving optimum, (*m*) was reflecting the prior on the chronogram but not on the phylogram (Fig. 3A, blue violins). In summary, fitting an OU process on a phylogram results in biases not only for a trait under selection but also for a trait under neutral evolution.

## Discussion

In phylogenetic comparative methods, the evolution of continuous traits is typically modelled as a stochastic process running along the branches of a phylogenetic tree. The null model of neutral trait evolution, genetic drift, is thought of as a Brownian motion (BM) running on the tree (Lynch and Hill, 1986; Felsenstein, 1988), and deviations from this model are interpreted as selection acting on the trait (Butler and King, 2004). As a result, a trait on which the BM is favoured over alternative models are sometimes interpreted as a trait evolving neutrally (Khaitovich et al., 2006; Catalán et al., 2019). However, the BM is not only the null model of neutral evolution, but can also model selection, with a trait tracking a moving optimum, where the optimum itself is following a BM (Hansen and Martins, 1996; Latrille et al., 2024). Distinguishing these scenarios is not trivial, but we here showed that the underlying backbone tree to model trait evolution can help disentangle neutral evolution from selection. The standard in phylogenetic comparative methods is a chronogram, where the branch lengths are measured in time (Felsenstein, 1985; Harmon, 2018), while phylograms, where the branch lengths are measured in number of substitutions instead, have been generally overlooked. For a simulated trait under selection we assessed that the BM has indeed a better fit on a chronogram than on a phylogram. However, we argue that to model a trait evolving neutrally, the backbone tree should not be a chronogram but instead a phylogram. Phylograms better represent the number of generations that occurred, and in turn should explain best the differences in traits between species (Latrille et al., 2024) In practice, we show that for simulations of a trait evolving neutrally, the fit of a BM supports a phylogram rather than a chronogram. Additionally, for a neutral trait, both ancestral state reconstruction and estimation for the number of rate changes for a multi-rate BM was more accurate on a phylogram. Altogether, the combined use of a phylogram and a chronogram can help support the hypothesis of neutral evolution, or to rule it out.

We applied this method to different empirical datasets of body and brain masses in mammals, and we showed that the chronogram was favoured over the phylogram, ruling out the neutral model of evolution for these traits (Latrille et al., 2024). The mammalian dataset is particularly relevant, since changes in generation time are expected to occur along the phylogenetic tree, from 6 months for mice to 50 years for bowhead whales (De Magalhães and Costa, 2009; Nielsen et al., 2016). But more generally, our method is also relevant when changes in mutation rate (per generation) and effective population size (*N*_e_) are expected. First, normalizing by nucleotide divergence also accounts theoretically for variations in effective population size (*N*_e_) since the substitution rate of neutral mutations is equal to the mutation rate, such that *N*_e_ has no effect on the rate of neutral evolution (Kimura, 1968; Ohta, 1972). Moreover, if population structure were to impact the probability of fixation for neutral mutations (which would also impact a neutral trait), the phylogram would automatically absorb these changes. This argument was partially tested in our simulation by modelling changes in both *N*_e_ and generation time (although not modelling population structure *per se*), and the phylogram was favoured over the chronogram for a trait evolving neutrally. Second, under certain conditions, the use of phylograms can also absorb changes in mutation rate (per loci per generation), which in mammals can vary up to 10 fold (Bergeron et al., 2023). As in the case of our simulations, we assumed that the genetic architecture of the trait is constant and the mutation rate is homogeneous across the genome. However, if the mutation rate would only change for a subset of the genome that is encoding the trait, the phylogram would not absorb these changes in mutation rate. Finally, because chronograms are usually derived from phylograms by calibrating the tree with molecular clocks, using phylograms directly has the additional advantage of removing model assumptions required to calibrate node ages (Litsios and Salamin, 2012).

These examples show that while phylograms are useful to model neutral evolution, they are not a silver bullet to disentangle the effect of drift and selection. In our simulations, we generally found that using a phylogram tends to favour the OU process over BM. This suggests that trees in units of nucleotide divergence may not always provide a reliable framework for modelling trait evolution, especially when selection is involved. We recommend continuing to use chronograms for modelling selection acting on a trait, as they generally provide more accurate and reliable results. Moreover, for a trait under selection for which the pace of optimum changes is also tracking the changes in generation time, the support for a phylogram over a chronogram would be misleading (Litsios and Salamin, 2012). Therefore, we argue that while phylograms can offer valuable insights (Wilson et al., 2022), they should be used with caution and in conjunction with chronograms to provide a more comprehensive understanding of trait evolution. Contrarily to our cautious note, Wilson et al. (2022) suggested that phylograms are more robust for studying discrete character evolution than chronogram. This apparent contradiction is due to the different units of measurement used in the studies, as they measure branch lengths in units of morphological distance rather than substitutions per site (Wilson et al., 2022). We thus argue that the use of a phylogram, measured in units of substitutions, could also be useful for studying the evolution of discrete characters, but the same caution should be applied.

Caution is indeed warranted, as there are several limitations to our approach. First, we simulated only a subset of different regimes of selection, but more complex scenarios could be included, for example with primary and secondary moving optima (Hansen, 2024). Additionally, the simulated genetic architecture of a single trait is assumed to be constant across the phylogeny. Constant architecture results in the saturation of the phenotype on long time scales, with a phenomenon similar to nucleotide saturation (Latrille et al., 2024). This saturation effect might be one of the reasons for a bias in favour of the OU process over BM that we observed in our simulations. A more realistic genetic architecture should include pleiotropy, epistasis and linkage disequilibrium, while here we only considered non-linked loci contributing additively to a single trait. Second, while the tested alternative models of evolution (multi-rates BM, OU, multi-optimum OU) are standard, they do not encompass the full breadth of available methods to model trait evolution (Pennell et al., 2014; Höhna et al., 2016). Finally, the mammalian dataset (brain and body) may not be representative of all traits or species and even though our analysis showcases the utility of phylograms, more empirical studies are needed to replicate the findings.

One striking example of a trait that could (and should) be empirically studied with a phylogram is gene expression level, both for mRNA and protein levels. Indeed, the regime of selection acting on gene expression level is the focus of intense debate (Signor and Nuzhdin, 2018; Price et al., 2022; Bertram et al., 2023; Dimayacyac et al., 2023). Typically, datasets on which BM is favoured over other models is interpreted as mRNA expression level evolving neutrally (Khaitovich et al., 2006; Catalán et al., 2019; Dimayacyac et al., 2023). Phenotypic distance is also plotted as a function of time while claiming that a trait is evolving under drift (Jiang et al., 2023). Instead, we argue that phenotypic distance should be shown as a function of branch length in units of substitutions (square root), not time. More generally, from an empirical perspective, the use of nucleotide changes at neutral loci as a normalizing factor brings new perspectives into trait evolution (Latrille et al., 2024).

Altogether, our study supports the use of a chronogram when testing for selection acting on a trait. The pipeline of hypothesis testing and model comparison is currently well established and should be followed. However, when the analysis concludes that the BM is favoured, then the support of the phylogram over the chronogram should be tested additionally. We provide a Bayesian model implemented in *RevBayes* to test this hypothesis, which indicates whether a BM supports better a phylogram or a chronogram (0 ≤ *π* ≤ 1), given the data at the tip of the tree and both trees with the same topology. Only if the phylogram has more support than the chronogram (*π* close to 1) can it be assumed that the trait is evolving under drift, but caution is still warranted.

## Author contributions

Original idea: T.L. and anonymous reviewers; Model conception: T.L., T.G. and N.S.; Code: T.L.; Data analyses: T.L. and T.G.; Interpretation: T.L., T.G. and N.S.; First draft: T.L.; Editing and revisions: T.L., T.G. and N.S. Project management and funding: N.S.

## Acknowledgements

We gratefully acknowledge the help of Lucy M. Fitzgerald, Diego A. Hartasánchez, Anna Marcionetti and Julien Joseph for their advice and reviews concerning this manuscript.

## Data and materials availability

The materials that support the findings of this study are openly available in GitHub at github.com/ThibaultLatrille/ChronoPhylogram. Snakemake pipeline, simulator, analysis scripts and *RevBayes* scripts and documentation are available in the repository to replicate the study.

## References

Barton, N. H., A. M. Etheridge, and A. Véber (2017). “The Infinitesimal Model: Definition, Derivation, and Implications”. In: Theoretical Population Biology 118, pp. 50–73.

Beaulieu, J. M., D.-C. Jhwueng, C. Boettiger, and B.C. O’Meara (2012). “Modeling Stabilizing Selection: Expanding the Ornstein-Uhlenbeck Model of Adaptive Evolution”. In: Evolution 66.8, pp. 2369–2383.

Bergeron, L. A., S. Besenbacher, J. Zheng, P. Li, M. F. Bertelsen, B. Quintard, J. I. Hoffman, Z. Li, J. St. Leger, C. Shao, J. Stiller, M. T. P. Gilbert, M. H. Schierup, and G. Zhang (2023). “Evolution of the Germline Mutation Rate across Vertebrates”. In: Nature, pp. 1–7.

Bertram, J., B. Fulton, J. P. Tourigny, Y. Peña-Garcia, L. C. Moyle, and M. W. Hahn (2023). “CAGEE: Computational Analysis of Gene Expression Evolution”. In: Molecular Biology and Evolution 40.5, msad106

Blanquart, F. and T. Bataillon (2016). “Epistasis and the Structure of Fitness Landscapes: Are Experimental Fitness Landscapes Compatible with Fisher’s Geometric Model?” In: Genetics 203.2, pp. 847–862.

Butler, M. A. and A. A. King (2004). “Phylogenetic Comparative Analysis: A Modeling Approach for Adaptive Evolution.” In: The American naturalist 164.6, pp. 683–695.

Catalán, A., A. D. Briscoe, and S. Höhna (2019). “Drift and Directional Selection Are the Evolutionary Forces Driving Gene Expression Divergence in Eye and Brain Tissue of Heliconius Butterflies”. In: Genetics 213.2, pp. 581–594.

Cooper, N., G. H. Thomas, C. Venditti, A. Meade, and R. P. Freckleton (2016). “A Cautionary Note on the Use of Ornstein Uhlenbeck Models in Macroevolutionary Studies”. In: Biological Journal of the Linnean Society 118.1, pp. 64–77.

De Magalhães, J. P. and J. Costa (2009). “A Database of Vertebrate Longevity Records and Their Relation to Other Life-history Traits”. In: Journal of Evolutionary Biology 22.8, pp. 1770–1774.

Dimayacyac, J. R., S. Wu, D. Jiang, and M. Pennell (2023). “Evaluating the Performance of Widely Used Phylogenetic Models for Gene Expression Evolution”. In: Genome Biology and Evolution, evad211.

Eastman, J. M., M. E. Alfaro, P. Joyce, A. L. Hipp, and L. J. Harmon (2011). “A Novel Comparative Method for Identifying Shifts in the Rate of Character Evolution on Trees”. In: Evolution 65.12, pp. 3578–3589.

Felsenstein, J. (1985). “Phylogenies and the Comparative Method”. In: The American Naturalist 125.1, pp. 1–15.

(1988). “Phylogenies and Quantitative Characters”. In: Annual Review of Ecology and Systematics 19.1, pp. 445–471.

Foley, N. M., V. C. Mason, A. J. Harris, K. R. Bredemeyer, J. Damas, H. A. Lewin, E. Eizirik, J. Gatesy, E. K. Karlsson, K. Lindblad-Toh, Zoonomia Consortium, M. S. Springer, and W. J. Murphy (2023). “A Genomic Timescale for Placental Mammal Evolution”. In: Science 380.6643, eabl8189.

Genereux, D. P. et al. (2020). “A Comparative Genomics Multitool for Scientific Discovery and Conservation”. In: Nature 587.7833, pp. 240–245.

Grabowski, M., J. Pienaar, K. L. Voje, S. Andersson, J. Fuentes-González, B. T. Kopperud, D. S. Moen, M. Tsuboi, J. Uyeda, and T. F. Hansen (2023). “A Cautionary Note on “A Cautionary Note on the Use of Ornstein Uhlenbeck Models in Macroevolutionary Studies”“. In: Systematic Biology 72.4, pp. 955–963.

Hansen, T. F. (2024). “Three Modes of Evolution?Remarks on Rates of Evolution and Time Scaling”. In: Journal of Evolutionary Biology, voae071.

(1997). “Stabilizing Selection and the Comparative Analysis of Adaptation”. In: Evolution 51.5, pp. 1341– 1351.

Hansen, T. F. and E. P. Martins (1996). “Translating between Microevolutionary Process and Macroevolutionary Patterns: The Correlation Structure of Interspecific Data”. In: Evolution 50.4, pp. 1404–1417.

Hansen, T. F., J. Pienaar, and S. H. Orzack (2008). “A Comparative Method for Studying Adaptation to a Randomly Evolving Environment”. In: Evolution 62.8, pp. 1965–1977.

Harmon, L. (2018). “Phylogenetic Comparative Methods: Learning from Trees”. In.

Höhna, S., M. J. Landis, T. A. Heath, B. Boussau, N. Lartillot, B. R. Moore, J. P. Huelsenbeck, and F. Ronquist (2016). “RevBayes: Bayesian Phylogenetic Inference Using Graphical Models and an Interactive Model-Specification Language”. In: Systematic Biology 65.4, pp. 726–736.

Holstad, A., K. L. Voje, Ø. H. Opedal, G. H. Bolstad, S. Bourg, T. F. Hansen, and C. Pélabon (2024). “Evolvability Predicts Macroevolution under Fluctuating Selection”. In: Science 384.6696, pp. 688–693.

Ingram, T. and D. Mahler (2013). “SURFACE: Detecting Convergent Evolution from Comparative Data by Fitting Ornstein-Uhlenbeck Models with Stepwise Akaike Information Criterion”. In: Methods in Ecology and Evolution 4.5, pp. 416–425.

Jiang, D., A. L. Cope, J. Zhang, and M. Pennell (2023). “On the Decoupling of Evolutionary Changes in mRNA and Protein Levels”. In: Molecular Biology and Evolution 40.8, msad169.

Khabbazian, M., R. Kriebel, K. Rohe, and C. Ané (2016). “Fast and Accurate Detection of Evolutionary Shifts in Ornstein–Uhlenbeck Models”. In: Methods in Ecology and Evolution 7.7, pp. 811–824.

Khaitovich, P., W. Enard, M. Lachmann, and S. Pääbo (2006). “Evolution of Primate Gene Expression”. In: Nature Reviews Genetics 7.9, pp. 693–702.

Kimura, M. (1968). “Evolutionary Rate at the Molecular Level”. In: Nature 217.5129, pp. 624–626.

(1983). The Neutral Theory of Molecular Evolution. Cambridge University Press.

Lande, R. (1976). “Natural Selection and Random Genetic Drift in Phenotypic Evolution”. In: Evolution 30.2, pp. 314–334. JSTOR: 2407703.

(1980). “The Genetic Covariance between Characters Maintained by Pleiotropic Mutations”. In: Genetics 94.1, pp. 203–215.

Latrille, T., M. Bastian, T. Gaboriau, and N. Salamin (2024). “Detecting Diversifying Selection for a Trait from within and Between-Species Genotypes and Phenotypes”. In: Journal of Evolutionary Biology 37.12, pp. 1538–1550.

Litsios, G. and N. Salamin (2012). “Effects of Phylogenetic Signal on Ancestral State Reconstruction”. In: Systematic Biology 61.3, pp. 533–538.

Lynch, M. and W. G. Hill (1986). “Phenotypic Evolution by Neutral Mutation”. In: Evolution 40.5, pp. 915– 935.

Mitov, V., K. Bartoszek, G. Asimomitis, and T. Stadler (2020). “Fast Likelihood Calculation for Multivariate Gaussian Phylogenetic Models with Shifts”. In: Theoretical Population Biology 131, pp. 66–78.

Nielsen, J., R. B. Hedeholm, J. Heinemeier, P. G. Bushnell, J. S. Christiansen, J. Olsen, C. B. Ramsey, R. W. Brill, M. Simon, K. F. Steffensen, and J. F. Steffensen (2016). “Eye Lens Radiocarbon Reveals Centuries of Longevity in the Greenland Shark (Somniosus Microcephalus)”. In: Science 353.6300, pp. 702–704.

Ohta, T. (1972). “Population Size and Rate of Evolution”. In: Journal of Molecular Evolution 1.4, pp. 305– 314.

Paradis, E., J. Claude, and K. Strimmer (2004). “APE: Analyses of Phylogenetics and Evolution in R Language”. In: Bioinformatics 20.2, pp. 289–290.

Pennell, M. W., J. M. Eastman, G. J. Slater, J. W. Brown, J. C. Uyeda, R. G. FitzJohn, M. E. Alfaro, and L. J. Harmon (2014). “Geiger v2.0: An Expanded Suite of Methods for Fitting Macroevolutionary Models to Phylogenetic Trees”. In: Bioinformatics 30.15, pp. 2216–2218.

Price, P. D., D. H. Palmer Droguett, J. A. Taylor, D. W. Kim, E. S. Place, T. F. Rogers, J. E. Mank, C. R. Cooney, and A. E. Wright (2022). “Detecting Signatures of Selection on Gene Expression”. In: Nature Ecology & Evolution, pp. 1–11.

Sella, G. and N. H. Barton (2019). “Thinking about the Evolution of Complex Traits in the Era of GenomeWide Association Studies”. In: Annual Review of Genomics and Human Genetics 20.1, pp. 461–493.

Signor, S. A. and S. V. Nuzhdin (2018). “The Evolution of Gene Expression in Cis and Trans”. In: Trends in Genetics 34.7, pp. 532–544.

Silvestro, D., A. Kostikova, G. Litsios, P. B. Pearman, and N. Salamin (2015). “Measurement Errors Should Always Be Incorporated in Phylogenetic Comparative Analysis”. In: Methods in Ecology and Evolution 6.3, pp. 340–346.

Silvestro, D., M. F. Tejedor, M. L. Serrano-Serrano, O. Loiseau, V. Rossier, J. Rolland, A. Zizka, S. Höhna, A. Antonelli, and N. Salamin (2019). “Early Arrival and Climatically-Linked Geographic Expansion of New World Monkeys from Tiny African Ancestors”. In: Systematic Biology 68.1, pp. 78–92.

Soria, C. D., M. Pacifici, M. Di Marco, S. M. Stephen, and C. Rondinini (2021). “COMBINE: A Coalesced Mammal Database of Intrinsic and Extrinsic Traits”. In: Ecology 102.6, e03344.

Tenaillon, O. (2014). “The Utility of Fisher’s Geometric Model in Evolutionary Genetics”. In: Annual Review of Ecology, Evolution, and Systematics 45.1, pp. 179–201.

Tsuboi, M., W. van der Bijl, B. T. Kopperud, J. Erritzøe, K. L. Voje, A. Kotrschal, K. E. Yopak, S. P. Collin, A. N. Iwaniuk, and N. Kolm (2018). “Breakdown of Brain–Body Allometry and the Encephalization of Birds and Mammals”. In: Nature Ecology & Evolution 2.9, pp. 1492–1500.

Turelli, M. (1984). “Heritable Genetic Variation via Mutation-Selection Balance: Lerch’s Zeta Meets the Abdominal Bristle”. In: Theoretical Population Biology 25.2, pp. 138–193.

Uyeda, J. C. and L. J. Harmon (2014). “A Novel Bayesian Method for Inferring and Interpreting the Dynamics of Adaptive Landscapes from Phylogenetic Comparative Data”. In: Systematic Biology 63.6, pp. 902–918.

Wilson, J. D., N. Mongiardino Koch, and M.J. Ramírez (2022). “Chronogram or Phylogram for Ancestral State Estimation? Model-fit Statistics Indicate the Branch Lengths Underlying a Binary Character’s Evolution”. In: Methods in Ecology and Evolution 13.8, pp. 1679–1689.

