## Supplementary materials for "Phylogram instead of chronogram when assessing the neutral evolution of a trait"

#### Contents

1 Simulation of continuous trait 16
1.1 Genotype-phenotype map . . . . . . . . . . . . . . . . . . . . . . . . . . . . . . . . 16
1.2 Simulation along a tree . . . . . . . . . . . . . . . . . . . . . . . . . . . . . . . . . 16
2 Ancestral trait reconstruction 18
3 Empirical dataset 19

#### 1 Simulation of continuous trait

##### 1.1 Genotype-phenotype map

- $L$ is the number of loci encoding the trait.
- $a_l \sim \mathcal{N}(0, a^2)$ is the effect of a mutation on the trait at locus $l \in \{1, \dots, L\}$.
- $N_\text{e}$ is the effective number of individuals.
- $g_{k,l} \in \{0, 1, 2\}$ is the genotypic value at locus $l$ for individual $k \in \{1, \dots, N_\text{e}\}$.
- $G_k = \sum_{l=1}^{L} a_l \times g_{k,l}$ is the genotypic value for individual $k$.
- $\xi_k \sim \mathcal{N}(0, V_\text{E})$ is the effect of environment on the trait for individual $k$.
- $P_k = G_k + \xi_i$ is the phenotype for individual $k$.

Figure S1: summary of trait's genetic architecture.

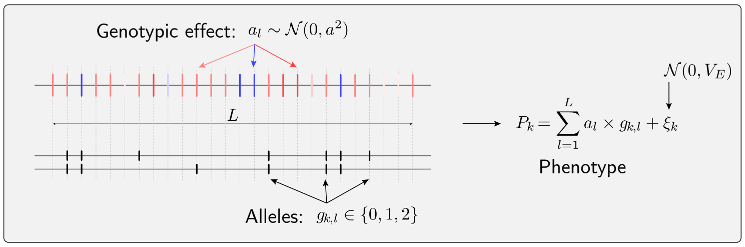


##### 1.2 Simulation along a tree

Each individual phenotypic value was the sum of genotypic value and an environmental effect. The environmental effect was normally distributed with variance $V_\text{E}$. We assumed that the genotypic value was encoded by $L = 5,000$ loci, with each locus contributing an additive effect that was normally distributed with standard deviation $a = 1$. We assumed a trait with a narrow-sense heritability of $h^2 = 0.2$ and computed the theoretical $V_\text{E}$ accordingly. Assuming a diploid panmictic population of size $N_\text{e} = 50$ at the root of the tree, and with non-overlapping generations, we simulated explicitly each generation along an ultrametric phylogenetic tree. For each offspring, the number of mutations was drawn from a Poisson distribution with mean $2 \cdot \mu \cdot L$, with the mutation rate per locus per generation $\mu$. From the empirical mammalian dataset (see Methods), we computed an average nucleotide divergence from the root to leaves of 0.18. We scaled parameters in our simulations to fit plausible values for mammals. We thus used a nucleotide mutation rate of $u = 0.00276/4N_{\mathrm{e}} = 1.38 \times 10^{-5}$ per site per generation and a total of $0.18/1.38 \times 10^{-5} = 13,500$ generations from root to leaves, and the number of generations along each branch was proportional to the branch length. We set $\mu = u$ without loss in generality since the genetic architecture ($L$ and $a$) is assumed constant in the simulator.

The changes in $\mu$ and $N_{\mathrm{e}}$ along the lineages were both modelled by a Brownian motion (BM) on the log scale (log-$\mu$ and log-$N_{\mathrm{e}}$), leading to geometric Brownian motion on the linear scale ($\mu$ and $N_{\mathrm{e}}$). These processes are parametrized as $\mathcal{B}\left(0, \sigma_{\mu} = 0.0043\right)$ and $\mathcal{B}\left(0, \sigma_{N_{\mathrm{e}}} = 0.0043\right)$, which, if counted across 13,500 generations, leads to a standard deviation of $0.0043 \cdot \sqrt{13,500} = 0.5$. In other words, the deviation in log-$N_{\mathrm{e}}$ and log-$\mu$ between the extant species and the root is 0.5. An Ornstein-Uhlenbeck process was overlaid to the instant value of log-$N_{\mathrm{e}}$ provided by the geometric BM to account for short-term changes between generations $\left(\mathrm{OU}\left(0, \sigma_{N_{\mathrm{e}}} = 0.1, \theta_{N_{\mathrm{e}}} = 0.9\right)\right)$. The geometric Brownian motion accounted for long-term fluctuations (low rate of changes $\sigma_{N_{\mathrm{e}}}$ but unbounded), while the Ornstein-Uhlenbeck introduced short-term fluctuations (high rate of changes $\sigma_{N_{\mathrm{e}}}$ but bounded and mean-reverting). The simulation started from an initial sequence at equilibrium at the root of the tree and, at each node, the process was split until it finally reached the leaves of the tree. From a speciation process perspective, this was equivalent to an allopatric speciation over one generation.

At each generation, parents were randomly sampled with a weight proportional to their fitness ($W$). Selection was modelled as a one-dimensional Fisher's geometric landscape, with the fitness of an individual being a monotonously decreasing function of the distance between the individual and the optimal phenotype (Tenaillon, 2014; Blanquart and Bataillon, 2016). More specifically, the fitness of an individual was given by $W = \mathrm{e}^{(P-\lambda)^2/\alpha}$, where $P$ was the trait value of the individual, $\lambda = 0.0$ was the optimal trait value, and $\alpha = 0.02$ was the strength of selection. Mutations were considered as a displacement of the phenotype in the multidimensional space. Beneficial mutations moved the phenotype closer to the optimum, while deleterious mutations moved it further away. Moving optimum selection was implemented by allowing the optimum phenotype to move along the phylogenetic tree as a geometric BM (Hansen, 1997) $\left(\lambda \sim \mathcal{B}\left(0, \sigma_{\lambda} = 1.0\right)\right)$. Multiple optima were implemented by allowing the optimum phenotype to shift along a branch with a probability of $10^{-5}$, the shift in optimum being then drawn from a exponential distribution with rate parameter 0.1, meaning an average of 10 in the optimum shift per switch. Neutral evolution was implemented by flattening the fitness landscape ($W = 1$), which meant that each individual had the same probability of being sampled at each generation.

#### 2 Ancestral trait reconstruction

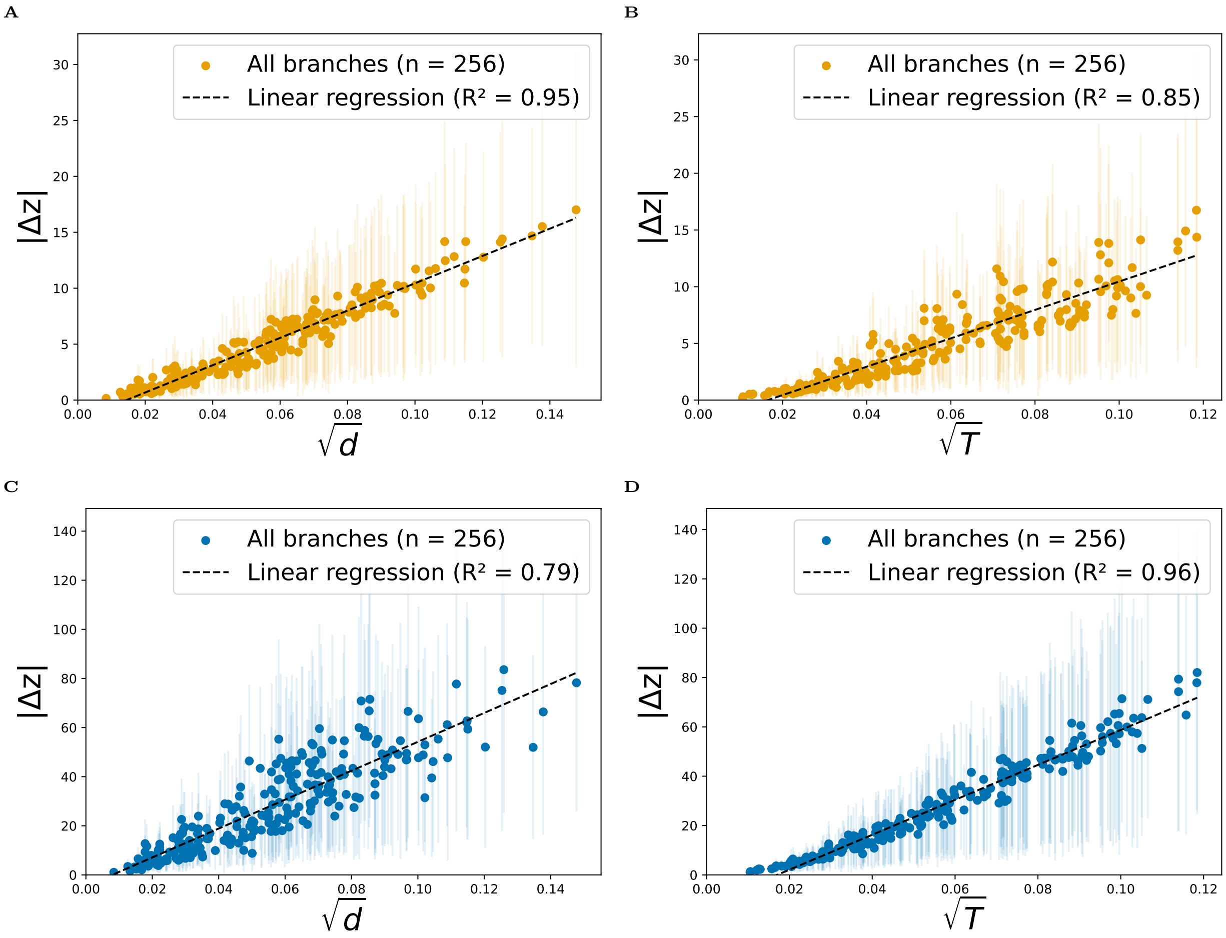


Figure S2: **Phenotypic distance as a function of branch length.** Phenotypic distance ($|\Delta z|$) is computed as the absolute difference of the posterior mean trait value between the ancestral node and the descendent node. Across all simulated dataset (100 replicates), we computed the mean phenotypic distance (blue dots) and standard deviation (blue horizontal lines) for each branch. Simulations under a neutral regime (yellow, top row, panels A and B) and under a moving optimum regime (bottom row, panels C and D). Reconstruction of ancestral trait on a phylogram (blue, left column, panels A and C) and a chronogram (right column, panels B and D).

### 3 Empirical dataset

| Dataset | Trait | Sex | Likelihood | Taxa | Phylogram support ($\pi$) |
|---|---|---|---|---|---|
| *Mammal* | Body mass | Mixed | Full | 125 | 0.0 |
| *Mammal* | Body mass | Mixed | REML | 125 | 0.002 |
| *Mammal* | Body mass | ♀ | Full | 46 | 0.848 |
| *Mammal* | Body mass | ♀ | REML | 46 | 0.846 |
| *Mammal* | Body mass | ♂ | Full | 46 | 0.697 |
| *Mammal* | Body mass | ♂ | REML | 46 | 0.684 |
| *Mammal* | Brain mass | Mixed | Full | 125 | 0.0035 |
| *Mammal* | Brain mass | Mixed | REML | 125 | 0.0035 |
| *Mammal* | Brain mass | ♀ | Full | 46 | 0.56 |
| *Mammal* | Brain mass | ♀ | REML | 46 | 0.539 |
| *Mammal* | Brain mass | ♂ | Full | 46 | 0.515 |
| *Mammal* | Brain mass | ♂ | REML | 46 | 0.497 |
| *COMBINE* | Body mass | Mixed | Full | 217 | 0.0 |
| *COMBINE* | Body mass | Mixed | REML | 217 | 0.0 |
| *COMBINE* | Brain mass | Mixed | Full | 174 | 0.0015 |
| *COMBINE* | Brain mass | Mixed | REML | 174 | 0.00649 |

Table S1: **Summary of empirical datasets.** Support for the phylogram over the chronogram ($\pi$) for different datasets and traits. Brain and body mass is either extracted from Tsuboi et al. (2018, *Mammal*) or from the *COMBINE* database (Soria et al., 2021). For the Tsuboi et al. (2018, *Mammal*) dataset, the individuals can also be split into males (♂) and females (♀). Likelihood is either computed with a full likelihood or with a restricted maximum likelihood (REML). The number of taxa is the number of species with data available for the trait.
